# LINKIN-associated proteins necessary for tissue integrity during collective cell migration

**DOI:** 10.1101/2023.02.08.527750

**Authors:** Chieh-Hsiang Tan, Kai-Wen Cheng, Heenam Park, Tsui-Fen Chou, Paul W. Sternberg

**Author notes:** for correspondence Emails: Chieh-Hsiang Tan, Paul W. Sternberg. denotes equal contribution (Chieh-Hsiang Tan, Kai-Wen Cheng).

## Abstract

Cell adhesion plays essential roles in almost every aspect of metazoan biology. LINKIN (Human: ITFG1, *Caenorhabditis elegans*: *lnkn-1*) is a conserved transmembrane protein that has been identified to be necessary for tissue integrity during migration. In *C. elegans*, loss of *lnkn-1* results in the detachment of the lead migratory cell from the rest of the developing male gonad. Previously, three interactors of ITFG1/*lnkn-1* – RUVBL1/*ruvb-1*, RUVBL2/*ruvb-2*, and alpha-tubulin – were identified by immunoprecipitation-mass spectrometry (IP-MS) analysis using human HEK293T cells and then validated in the nematode male gonad. The ITFG1-RUVBL1 interaction has since been independently validated in a breast cancer cell line model that also implicates the involvement of the pair in metastasis. Here, we showed that epitope-tagged ITFG1 localized to the cell surface of MDA-MB-231 breast cancer cells. Using IP-MS analysis, we identified a new list of potential interactors of ITFG1. *Loss-of-function* analysis of their *C. elegans* orthologs found that three of the interactors – ATP9A/*tat-5*, NME1/*ndk-1*, and ANAPC2/*apc-2* – displayed migratory detachment phenotypes similar to that of *lnkn-1*. Taken together with the other genes whose reduction-of-function phenotype is similar to that of *lnkn-1* (notably cohesion and condensin), suggests the involvement of membrane remodeling and chromosome biology in LINKIN-dependent cell adhesion and supports the hypothesis for a structural role of chromosomes in post-mitotic cells.

## Introduction

Cell adhesion is often necessary in biological processes that extend to more than one cell. Metazoan development and intercellular signal transduction require a fine-tuning of both the strength and the content of cell adhesion. As such, misregulation or alteration of cell adhesion proteins leads to a broad spectrum of pathological conditions (Hynes, 1999; Janiszewska et al., 2020). Abnormal cell adhesion is associated with cancer development. Tumor metastasis requires that malignant cells both acquire the ability to migrate and lose cell adhesion (Geiger and Peeper, 2009). Decades of research and the advancement of genetics and biochemistry has led to much knowledge of how cell adhesion is maintained, both in terms of mechanism as well as the molecules involved. However, key questions remained, such as how dynamic cell adhesion is maintained during cell migration (Collins and Nelson, 2015).

The gonad of *Caenorhabditis elegans* is shaped through a combination of elongation and a stereotypic migration guided by a migratory leader cell. In male worms, the leader cell is a somatic gonadal cell known as the linker cell (LC) (Hirsh et al., 1976; Kimble and Hirsh, 1979; Klass et al., 1976). At the leading edge of the elongating male gonad, the migratory linker cell leads a collective migration of a stalk of passive migratory cells. There are many similarities between this collective migration and some of the well-known examples described in other organisms (Scarpa and Mayor, 2016), including the specification of the leader cell through Notch signaling (Greenwald et al., 1983; Hellstrom et al., 2007; Ikeya and Hayashi, 1999; Llimargas, 1999; Suchting et al., 2007; Yochem et al., 1988). As with the leader cells in other collective migrations (Mayor and Etienne-Manneville, 2016), the linker cell is responsible for both determining the direction of the migration (Kimble and Hirsh, 1979) and for generating a significant part of the traction forces “dragging” the group in the migration (Kato et al., 2014). If detached from the rest of the gonad, the linker cell retains the ability to migrate while the rest of the gonad ceases to elongate (Chisholm, 1991; Kato et al., 2014; Sternberg and Horvitz, 1988). Maintaining the cell-to-cell adhesion under intense mechanical stress and, thus, the overall integrity of the migrating complex is all but essential for gonadogenesis. Therefore, the migrating male gonad in this transparent nematode is a useful genetic model for the study of cell adhesion dynamics during migration.

A few proteins have been identified to be essential for tissue integrity and the cell adhesiveness of the linker cell-led collective migration. These include cytoskeleton proteins, namely alpha-tubulin, TBA-2, and beta-tubulin, TBB-2; SMC proteins HIM-1, SMC-3, SMC-4; a homeobox transcription factor EGL-5; AAA+ ATPase superfamily proteins RUVB-1 and RUVB-2; and LNKN-1, a conserved but poorly characterized transmembrane protein (Chisholm, 1991; Kato et al., 2014; Schwarz et al., 2012). Based upon the expression profile of a similar cell (Hunt-Newbury et al., 2007), Kato et al. (2014) identified *lnkn-1* through the observation of a linker cell detachment phenotype. This was followed by a proteomic approach consisting of immunoprecipitation of the human *lnkn-1* ortholog ITFG1 and protein identification through mass spectrometry. The interactors of ITFG1 were then functionally assayed in the nematode model using RNA interference, thereby identifying TBA-2, TBB-2, RUVB-1, and RUVB-2 as sharing the highly specific linker cell detachment phenotype with LNKN-1 (Kato et al., 2014). The interaction between *lnkn-1*/ITFG1 and *ruvb-1*/RUVBL1 was later independently verified in a breast cancer cell line (Fan et al., 2017), which also associates the pair with breast cancer progression.

Here, we set out to identify new interactors of LNKN-1/ITFG1 to advance our understanding of its role in migratory cells. We started by identifying interactors of the human ITFG1 in the breast cancer cell line MDA-MB-231. The cell line selection was based on physiological/ pathological relevance and documented expression (Fan et al., 2017). Working with this same cell line, we validated the ITFG1-RUVBL1 interaction and showed that ITFG1 is localized to the cell surface, consistent with it being involved in cell adhesion. Through immunoprecipitation and mass spectrometry-based protein identification in ITFG1-expressing cells, 1756 and 373 potential interactors of ITFG1 were identified in ITFG1 transient and stably expressing cells, respectively. Of these, 180 were identified through two unbiased independent experiments. We then proceeded to test the findings in *C. elegans*, in worms carrying loss-of-function mutations in their orthologs. Analyzing phenotypes that were also observed in *lnkn-1* knock-out worms, we identified four other genes that resulted in worms with incomplete male gonad migrations: aminophospholipid translocase *tat-5*/ATP9A; nucleoside diphosphate kinase *ndk-1*/NME1; anaphase-promoting complex *apc-2*/ANAPC2; and metastasis-associated protein *lin-40*/MTA2. Of these, we found the migratory failure of worms carrying *tat-5*, *ndk-1*, or *apc-2* loss-of-function alleles is likely caused by the disassociation of the linker cell from the majority of the elongating gonad, similar to that of *lnkn-1*. Based on our findings and the genes that had been previously identified, we conclude that LINKIN-dependent cell adhesion likely involves membrane remodeling and chromosome biology.

## Results

### Analysis of ITFG1 interactions

We started by transiently expressing the full-length ITFG1 protein with a carboxy-terminal Myc tag in the MDA-MB-231 breast cancer cell line (Fig. S1). As expected, immunostaining using anti-ITFG1 antibodies shows that ITFG1 localized to the plasma membrane (Fig. 1A). To confirm the subcellular localization of ITFG1, we analyzed the subcellular fractionated cell lysates from ITFG1 overexpressing cells and found ITFG1 highly enriched in the membrane fraction, as detected at approximately 75 – 100 kDa (Fig. 1B, Fig. S1). The surface localization of ITFG1 in living cells was further confirmed by flow cytometry (Fig. 1C). The plasma membrane localization results are consistent with a previous study (Fan et al., 2017) of the same MDA-MB-231 breast cancer cell line as well our original study using HEK293T cells (Kato et al., 2014). The localization is also consistent with the subcellular localization of LNKN-1 in *C. elegans* (Kato et al., 2014).

**Fig 1.**
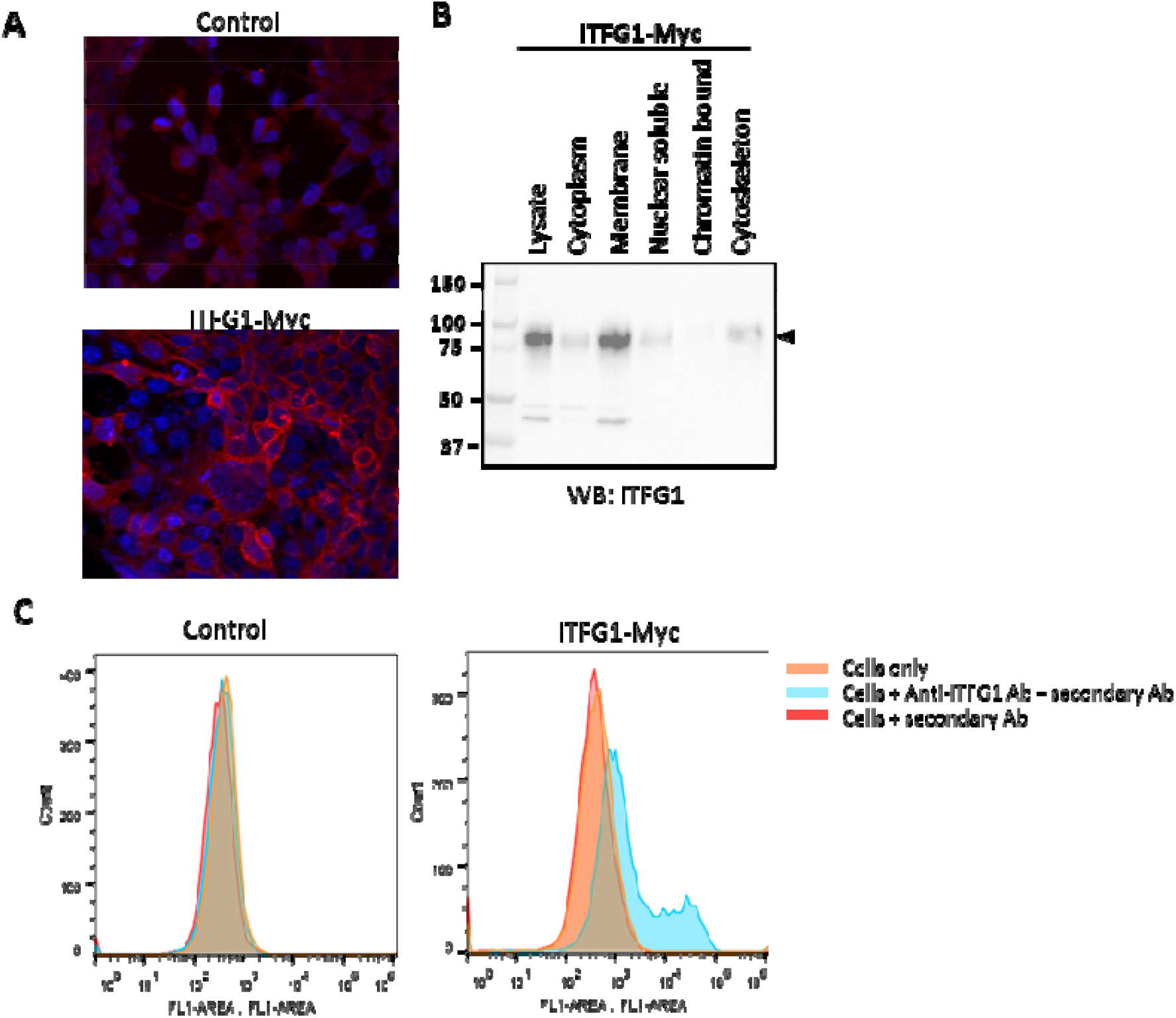
ITFG1 localized to the surface of MDA-MB-231 cells. (A) ITFG1 was detected on the plasma membrane in the ITFG1-Myc-expressing MDA-MB-231 cells using the anti-ITFG1 antibody (Red). Hoechst was used for nuclear staining (Blue). (B) ITFG1 was detected in the membrane fraction of the ITFG1 overexpressing cell lysates by western blot using anti-ITFG1 antibodies. (C) Flow cytometry confirms the surface localization of ITFG1 in living ITFG1-Myc-experessing MDA-MB-231 cells with anti-ITFG1 antibodies.

To identify other interactors of ITFG1, we also established a stable ITFG1-Myc-expressing MDA-MB-231 cells through lentivirus delivery. Cell lysates from both the transient and stable expression cultures were immunoprecipitated with anti-Myc agarose affinity gels and then analyzed by mass spectrometry. We identified a total of 2628 and 1242 proteins with higher enrichment (log2 FC > 0.6) in transient and stable ITFG1-expressing cells over control cells, respectively (Table S1-2). This mass spectrometry analysis identified both ITFG1 and its known interactors RUVBL1 and RUVBL2 (Fig. 2A, B), validating that our strategy of mass spectrometry coupled with immunoprecipitation as a method to identify ITFG1 interactors. To further validate the interactors identified with our approach, we also confirmed the ITFG1 and RUVBL1 interaction by showing that RUVBL1 can be reliably identified in the immunoprecipitated products of ITFG1-Myc-expressing cells and that ITFG1 can be reliably identified in the Flag-immunoprecipitated products of Flag-RUVBL1 expressing cells (Fig. S2). 180 proteins were enriched (log2 FC > 2) in both the immunoprecipitated products of transient and stable ITFG1-expressing cells (Table S3). Gene Ontology (GO) biological process and Reactome pathway enrichment analysis of these proteins found that the ITFG1 interacting proteins are enriched in multiple cell networks, including cell cycle, mitochondria translation initiation, and regulation of DNA repair (Fig. 2C).

**Fig 2.**
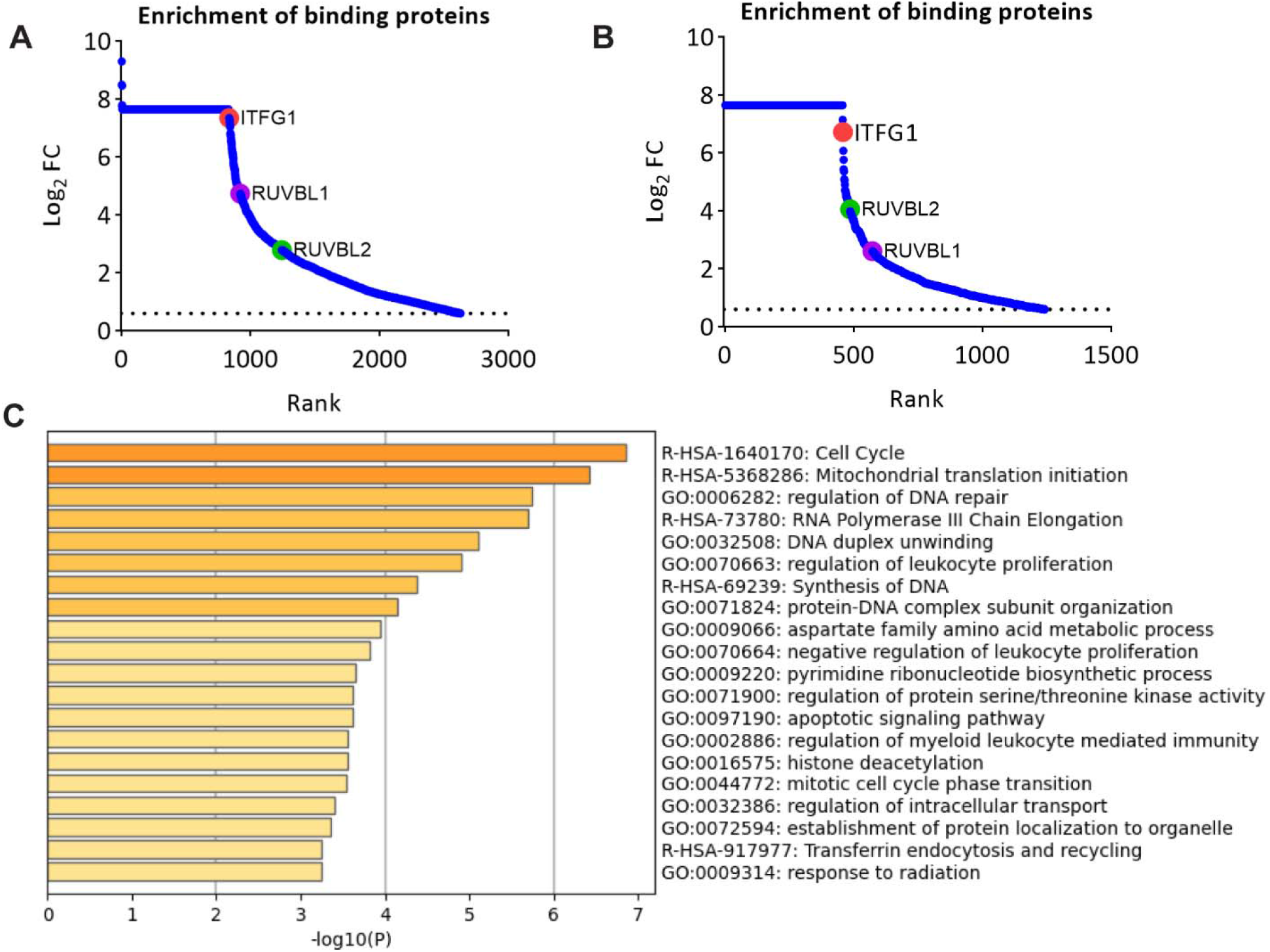
Identifying potential ITFG1 interactors with mass spectrometry. (A-B) fold change of the protein level in log2 between ITFG1-Myc-expressing cells over non-expressing control MDA-MB-231 cells. Each dot represents one protein. Red-ITFG1, Purple-RUVBL1, Green-RUVBL2. FC, fold change. (A) Transient ITFG1-Myc-expressing cells (B) Stable ITFG1-Myc-expressing cells. (C) Reactome pathway and GO biological process enrichment of the overlapping 180 proteins in ITFG-Myc expressing MDA-MB-231 cells. Data present the top 20 statistically enriched terms (*P*-adjusted value < 0.05).

### Mutations in the orthologs of ITFG1 interactors resulted in phenotypes similar to that of *lnkn-1* mutants

To validate the potential interactors and to explore their biological functions, we returned to the *C. elegans* model, where *lnkn-1* was originally identified (Kato et al., 2014). To select candidates for genetic analysis in this worm, we made our selection criteria more stringent, requiring candidates to be not only enriched in both sets of ITFG1 expressing cells but also detected in all the individual samples and at least >2.5 log2 fold enriched on average in the stable ITFG1 expressing cell line. Based on these stringent criteria, 84 *C. elegans* orthologs of 78 candidates were identified (Table S4), including *lnkn-1*/ITFG1 and the previously identified *ruvb-1*/RUVBL1 and *ruvb-2*/RUVBL2, as expected.

We reasoned that if the worm orthologs of the candidates interact with LNKN-1 in the cell adhesion process in *C. elegans*, loss-of-function of their encoding genes should display a phenotype similar to that of *lnkn-1(lf)*. Kato et al. (2014) reported multiple phenotypes associated with *lnkn-1* loss-of-function allele *gk367*, including that of the linker cell detachment. The allele *gk367*, generated by a *C. elegans* knock out consortium (Consortium, 2012), is a deletion that removes part of the *lnkn-1* coding sequence, presumably resulting in a truncated protein. Although the deletion only alters the coding sequence of *lnkn-1*, it is possible that *gk367* could also affect the expression of other genes co-transcribed in the same operon CEOP3552 (Davis et al., 2022). However, the phenotype observed in *gk367* animals is likely to resemble that of *lnkn-1(null)*, as the truncated protein is mislocalized (Kato et al., 2014), and knock-out of the next closest gene in the operon does not have similar effects (Tan et al., 2022). To generate a likely single gene knockout we used the STOP-IN cassette strategy (Wang et al., 2018) to generate *lnkn-1* allele *sy1596*. We used this allele for phenotypic analysis.

Hermaphrodites of both *lnkn-1(sy1596)* (Fig. 3) and *lnkn-1(gk256)* homozygous strains (Kato et al., 2014) are recessively maternal effect lethal; both *ruvb-1* and *ruvb-2* were also reported to be essential (Consortium, 2012; Updike and Mango, 2007). To select candidates for a comprehensive phenotypic analysis, we thus focused on genes that are likely to be essential. We reason that this is caused by the essentiality of a gene that is involved in the critical biological role of cell adhesion during development and that a significant amount of genetic information is provided maternally during nematode development (Evans and Hunter, 2005; Miwa et al., 1980; Wood et al., 1980). Therefore, we obtained deletion alleles for candidate genes based on their availability and whether the allele was phenotypically characterized as either lethal or sterile. To simplify developmental and fertility analysis, we sought to avoid the aneuploidy generated by translocation balancers. We thus rebalanced 19 lethal/sterile mutations primarily with the structurally defined and fluorescently and phenotypically labeled set of balancers described in Dejima et al. (2018) (Table S5). We then selected 17 candidate genes (Fig. 3C), starting with phenotypic analysis in the hermaphrodites. In all cases, homozygous animals descended from balanced heterozygous mothers were identified through the lack of fluorescently labeled balancers (Fig. 3A).

**Fig. 3.**
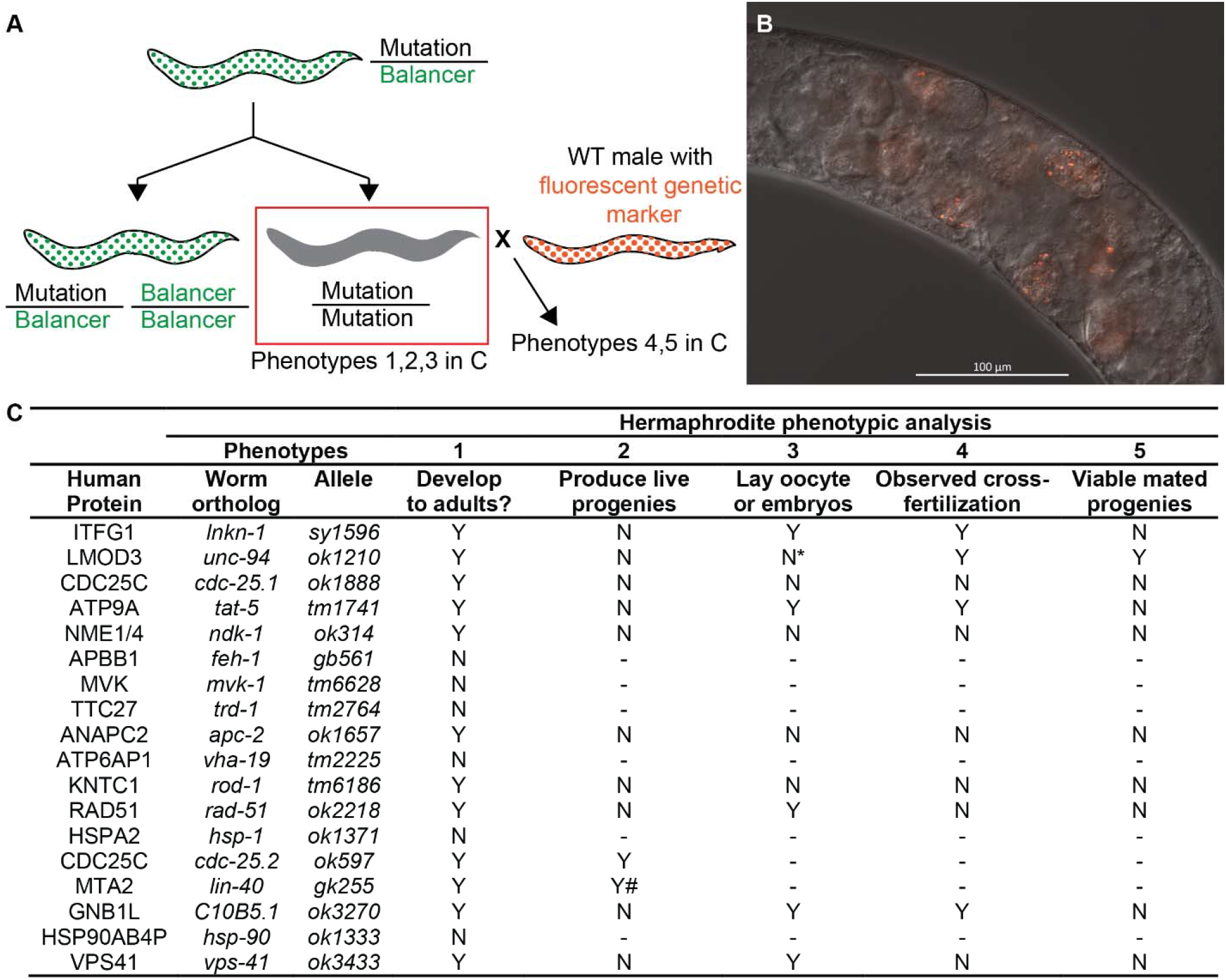
Mutations in the orthologs of ITFG1 interactors resulted in phenotypes similar to that of *lnkn-1* mutants. (A) Homozygotic animals of lethal/ sterile mutations used in this study descended from balanced heterozygotic mothers. Homozygotic animals were identified through the lack of fluorescently labeled balancers (green in the cartoon). In the maternal effect experiments, the homozygotic hermaphrodites were mated with wild-type males carrying a fluorescent genetic marker (*oxTi719* [*eft-3p::tdTomato::H2B*]) (Frokjaer-Jensen et al., 2014). (B) LNKN-1 is required maternally. Heterozygotic cross-progenies of *lnkn-1(sy1596)* hermaphrodites and *lnkn-1(+/+)* males fail to develop and die without hatching (The embryos carrying the orange-colored male-derived *oxTi719* marker can be seen). (C) Phenotypical analysis of the orthologs of ITFG1 interactors in *C. elegans*. Phenotype 2 and 5 was defined by having F_1_ progenies that hatch; Phenotype 3 was defined by embryos/ embryo-like objects or oocytes observed outside of the unmated hermaphrodite. Embryos and, in particular, oocytes may not be completely absent in all animals. This was especially the case in *unc-94(ok1210)*. Some *rod-1(tm6186)* animals also produce oocytes and embryo-like objects. Phenotype 4 was defined by the observation of F_1_ embryos/animals carrying the male-derived *oxTi719* marker. *: Very rarely; #: maternal effect sterile.

Mutants of seven candidate genes failed to develop into adults, arresting at various larval stages, preventing further characterization (Fig. 3C, phenotype 1). Of these, the arrest have been previously described for *feh-1(gb561)* (Zambrano et al., 2002), *trd-1(tm2764)* (Hughes et al., 2014), *hsp-1(ok1371)* (Knock Out Consortium), and *hsp-90(ok1333)* (Knock Out Consortium); but was previously unknown in *mvk-1(tm6628)*, *vha-19(tm2225)*, and *prx-3(tm6469)*. Although not pursued here, it is possible that these genes also contribute to LNKN-1-associated cell adhesion. The mutant animals from the remaining candidates were able to develop successfully into adults (Fig. 3C phenotype 1), and we next focused on fertility. As expected, most candidates failed to produce progeny that hatched, similar to *lnkn-1* loss-of-function mutants (Fig. 3C phenotype 2; (Kato et al., 2014). One major exception is *cdc-25.2(ok597)*, which we found to be fertile, consistent with a previous observation of low penetrance fertility (Kim et al., 2010). Another exception is *lin-40*, in which the *gk255* allele was selected for analysis based on the severity of the sterile phenotype we observed: *gk255* is maternal effect sterile while two other deletion alleles, *ok905* and *ok906,* were fertile.

We observed that unmated *lnkn-1(sy1596)* hermaphrodites produce a small number of oocytes and embryos that fail to hatch (Fig. 3C, phenotype 3). To test whether the embryonic contribution of LNKN-1 can alleviate this lethality, we crossed the *lnkn-1(sy1596)* hermaphrodites with wild-type males carrying a fluorescent genetic marker (Fig. 3A). The cross produced embryos with the male-derived fluorescent marker that still failed to hatch (Fig. 3B; Fig. 3C, phenotype 4,5), indicating that LNKN-1 is required in embryogenesis and needed to be provided maternally. Assessing the mutants of the candidate genes based on the *lnkn-1* phenotype mentioned above (Fig. 3C, phenotype 3,4,5), we find that *tat-5(tm1741)* and an uncharacterized gene *C10B5.1(ok3270)* are phenotypically similar to *lnkn-1*. *rad-51(ok2218)* and *vps-41(ok3433)* produced embryos that failed to hatch, and we were unable to obtain cross embryos through mating. On the other hand, although *unc-94(ok1210)* hermaphrodites rarely lay embryos or unfertilized oocytes by themselves (although they do produce them), it was able to produce mated heterozygous cross-progeny that hatch. This observation suggests that UNC-94 can be supplied both embryonically and maternally, unlike LNKN-1. No embryos or oocytes were observed outside of the hermaphrodites for *cdc-25.1(ok1888)*, *ndk-1(ok314)*, *apc-2(ok1657)*, and *rod-1(tm6186)*, and no mating resulted in cross progeny.

Hermaphrodite gonad development has a few key distinctions from that of the male: m: 1) In *C. elegans*, the hermaphrodite gonad is two-armed compared with one in the male and 2) the gonads of both males and hermaphrodites are tubes with a directional axis defined by the maturation of the germ cells, which occurs from distal to proximal (Kimble and Hirsh, 1979). However, the direction of gonad elongation is different: at the proximal end in males and the distal ends in hermaphrodites. 3) Partially as a consequence of the reversed direction of elongation, different cells serve the role of determining the migratory direction. On the proximal end in males, the linker cell (LC) serves as the leader cell in the migratory process. It both determines the direction of the migration and actively contributes to the process (Kato et al., 2014; Kimble and Hirsh, 1979). In hermaphrodites, with the primary elongation occurring at the other end of the tube, the linker cell analog – the anchor cell (AC) – is instead responsible for the induction of the vulva in the adjacent epidermal tissue (Kimble, 1981). Both cells are essential in their respective sexes in connecting the gonadal lumen to the exterior opening. In addition, in some other nematodes, the anchor cell also has a migratory role similar to the male linker cell (Felix and Sternberg, 1996). In hermaphrodites, it is the distal tip cells (DTC) that are responsible for determining the migratory direction (Kimble and White, 1981), playing a role similar to that of the linker cell in the males. However, as opposed to the linker cell, the distal tip cell does not in itself provide the traction for the migration, instead relying on the forces generated by the proliferating germ cells pushing it forward (Agarwal et al., 2022). Independent of their roles in migration, DTCs, two in the single-armed male gonad and one each in the two-armed hermaphrodite gonads, maintain the stem cell niche at the distal end, from which the germ cells proliferate (Kimble and White, 1981).

The gonads of *lnkn-1(sy1596)* hermaphrodites were noticeably different in shape from that of the wild-type (Fig. 4A, B), a phenotype previously noted in *lnkn-1(gk255)* (Kato et al., 2014). Each of the two U shaped gonad arms of the hermaphrodites can be roughly divided into two halves: the ventral-proximal half, which connects to the vulva and is shaped by the earlier centrifugal migration; and the dorsal-distal half, which is capped by the distal tip cell and is shaped by the later centripetal migration. In *lnkn-1(sy1596)* hermaphrodites, the centripetal half of the gonad is significantly shorter proportionally compared to that of the wild type (Fig. 4). This difference might be caused by either a late ventral to dorsal turn during the migration or a premature stop near the end of the elongation phase. We examined the nine sterile (Fig. 3 phenotype 2) candidates for similarities in morphology and found six of them, *cdc-25.1*, *tat-5*, *ndk-1*, *apc-2*, *rod-1*, and *C10B5.1,* share this phenotype. This phenotype has been previously reported in *ndk-1* and *apc-2* (Cram et al., 2006; Fancsalszky et al., 2014). Since the migration and elongation of the hermaphrodite gonad are driven by germ cell proliferation, and most if not all of the candidates produce much fewer embryos/ embryo-like objects compared to that of the wild-type, it is likely that at least part of the premature stopping could be due to reduced germ cell proliferation at later developmental stages. Indeed, one of the six candidates, namely *cdc-25.1,* is required for germ cell proliferation and described as having a reduced-sized gonad (Ashcroft and Golden, 2002; Kim et al., 2009; Yoon et al., 2012). It is worth noting that, similar to that of *lnkn-1*, all nine candidates have an overall reduced gonad size, but only mutants in six genes stop prematurely in the centripetal migration, so there are likely other factors involved.

**Fig. 4.**
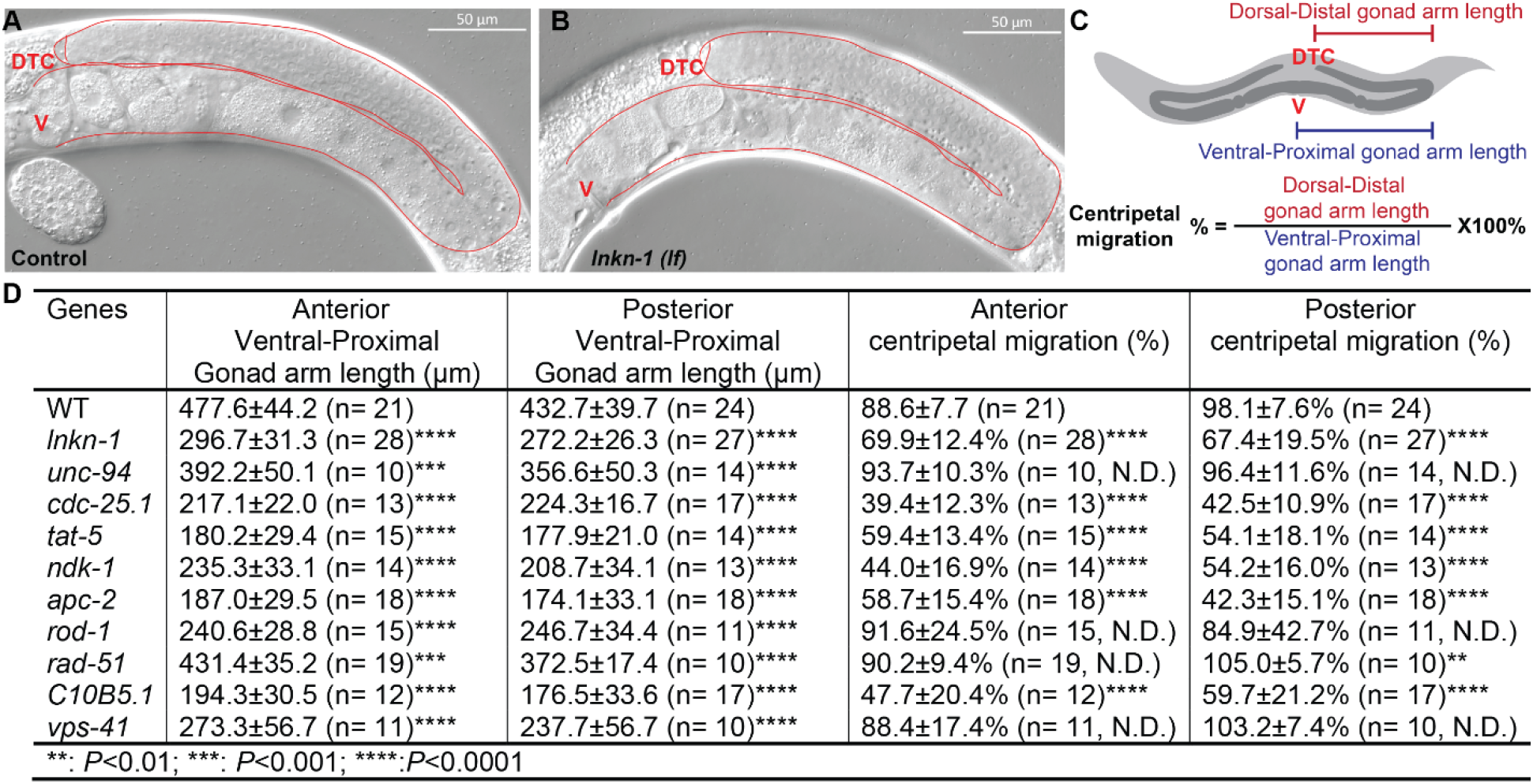
Potential LNKN-1 interactors displayed a similar gonad migration phenotype in hermaphrodites. (A-B). Loss of *lnkn-1* resulted in premature stopping of gonad migration in hermaphrodites (A) *lnkn-1(+)* control. (B) *lnkn-1(sy1596)* worms have a shortened dorsal-distal half of the gonad. (C-D) The premature stopping phenotype is defined by comparing the value of Centripetal migration (%) of the mutants to that of the wild-type (N2). The Centripetal migration (%) is determined by the ratio of dorsal-distal gonad arm length to that of ventral-proximal gonad arm length. The arm lengths were approximated as the distance from the vulval to the ventral-dorsal turn of the U-shaped gonad for the ventral-proximal gonad arm length; and the distance from the distal end of the gonad to the ventral-dorsal turn for the dorsal-distal gonad arm length. See materials and methods for detail. (D) Examination of gonad migration in hermaphrodites. Mutation in some of the candidate genes also causes pathfinding or other gonadal development defects in a number of animals. For the purpose of this examination, only gonads that made the centripetal migration was included. The pathfinding effects were consistently observed in *cdc-25.1(ok1888)* and *rod-1(tm6186)* and occurred at a lower frequency in *tat-5(tm1741)* and *apc-2(ok1657)* mutant animals. *Rod-1(tm6186)* animals were also observed to have missing gonad arms (n≥10. All comparisons were with the wild-type, **: p<0.01, ***: p<0.001, ****: p<0.0001, Student’s *t*-test).

### TAT-5/ATP9A, NDK-1/NME1, and APC-2/ANAPC2 are required for the integrity of the migrating male nematode gonad

We examined the mutant worms for a linker cell detachment phenotype. During the development of the *C. elegans* male, the linker cell leads a collective migratory process that shapes the male gonad and connects the gonadal lumen to the cloaca, opening up the passage for sperm (Kimble and Hirsh, 1979; Sulston et al., 1980). Disruption of this process, including the detachment of the linker cell, results in the male gonad not being connected to the exterior. We find that similar to that of *lnkn-1(lf)*, frequent occurrence of unconnected gonads were observed in male animals carrying mutant alleles of *tat-5*, *ndk-1*, *apc-2*, and *lin-40* (Fig. 5A).

**Fig. 5.**
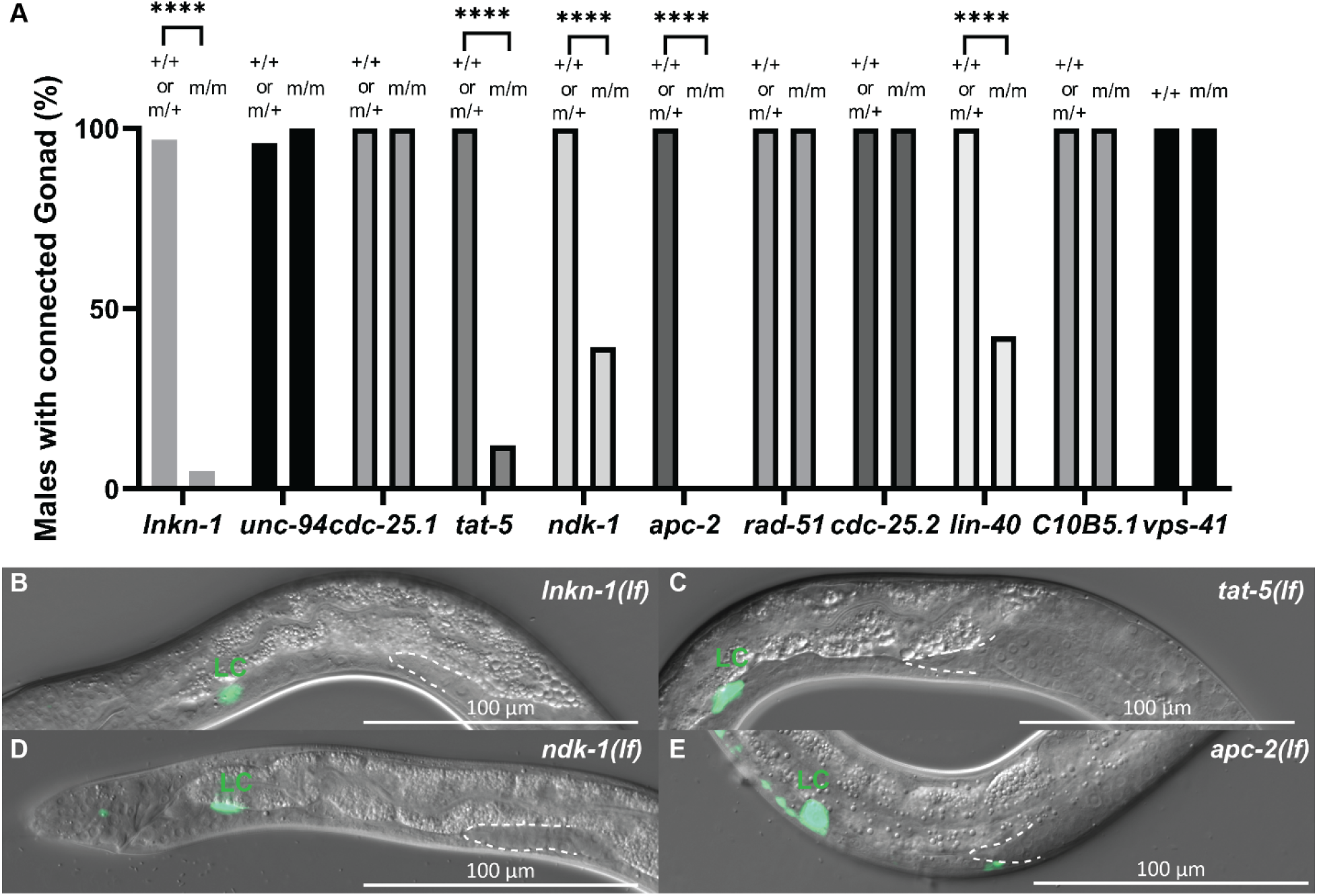
TAT-5/ATP9A, NDK-1/NME1, and APC-2/ANAPC2 are required for the integrity of the migrating male nematode gonad. (A) Males homozygotic for *tat-5(tm1741)*, *ndk-1(ok314)*, *apc-2(ok1657)*, and *lin-40(gk255)* have gonads that were not connected to the exterior similar to that of *lnkn-1(sy1596)*. For each genotype, the controls were siblings carrying the genetic balancer and were either the wild-type or heterozygous for the locus (n≥19, ****: p<0.0001, chi-square test). (B-E) Detached linker cells (LC) were found in males carrying *tat-5(tm1741)*, *ndk-1(ok314)*, or *apc-2(ok1657)*, similar to that of *lnkn-1(sy1596)*. Images are of animals in the L4 larvae stage, with *lag-2p::YFP* (syIs128) (colored in green) labeling the detached linker cell (LC). The posterior end of the remaining gonad is outlined in a white dashed line. (B) *lnkn-1(sy1596)*. (C) *tat-5(tm1741)*. (D) *ndk-1(ok314)*. (E) *apc-2(ok1657)*.

To determine whether the unconnected gonads were the results of linker cell detachment, we examined the gonads of developing animals. We found that as with *lnkn-1(lf)* (Fig. 5B), detached linker cells (LC) were observed in animals carrying the mutant alleles of *tat-5* (Fig. 5C)*, ndk-1* (Fig. 5D), and *apc-2* (Fig. 5E). The phenotype, however, was rarely observed in *lin-40(lf)*. Instead, the defect in *lin-40(lf)* animals seems to be caused by a late-stage pathfinding defect. Nonetheless, our results indicated that the *C. elegans* orthologs of ATP9A, NME1, NME4, and ANAPC2 are crucial for a physiological process that relies on cell adhesion. This observation and the fact that LNKN-1 is known to be critical for the same process give further credence to both their ITFG1 association and the role of LINKIN in cell adhesion.

## Discussion

By combining proteomics and association studies in cultured human cells with genetic analysis in *C. elegans*, we have identified three additional proteins that both are physically associated with LINKIN and play roles in the same process. This concordance of physical and genetic interaction places them firmly in the same cellular sub-network.

### Linker cell migration as a discovery platform for proteins involved of collective migration and adhesion

Some of the useful features of C. elegans – simplicity, invariant development, and a largely transparent body (Corsi et al., 2015) – apply strongly to the collective migration of the male gonad led by the linker cell. Starting in the L2 larvae stage, the linker cell migrated along a complicated trajectory that involves multiple turns, each occurring at a precise timing coordinating with the development of the animal, reaching the cloaca during L4 and underwent programmed cell death, opening up the sperm passage (Abraham et al., 2007; Hedgecock et al., 1987; Kato and Sternberg, 2009; Kimble and Hirsh, 1979). The entire process can be followed under microscopy, and the shape of the gonad can also be seen as a record of the path taken by the linker cell. While much more attention has been focused on the shaping of the hermaphroditic gonads, linker cells have been studied from a few different perspectives. First of all, it has been utilized as a model to study cell migration, often in conjunction with the study of other cell types, such as the axon migration in neurons and, in particular, the distal tip cell of the hermaphrodites (Antebi et al., 1998; Blelloch et al., 1999; Blelloch and Kimble, 1999; Clark et al., 1993; Hedgecock et al., 1990; Hedgecock et al., 1987; Kato et al., 2021; Kato and Sternberg, 2009; Nishiwaki, 1999; Nishiwaki et al., 2000; Palmer et al., 2002; Su et al., 2000; Tamai and Nishiwaki, 2007; Vogel and Hedgecock, 2001). The LC undergoes a caspase-independent form of programmed cell death (Abraham et al., 2007; Blum et al., 2012; Denning et al., 2013; Keil et al., 2017; Kinet et al., 2016; Lee et al., 2019; Malin et al., 2016; Schwendeman and Shaham, 2016). As part of the effort to understand its biology, the transcriptome of the cell, acquired through physical isolation, is known at different stages during its development (Schwarz et al., 2012). Finally, related to this study, linker cell has been studied in relation to cell adhesion (Kato et al., 2014).

In *C. elegans*, the male gonad development offers a unique opportunity to study postembryonic collective cell migration. While sharing many aspects in their development, particularly the migratory orientations, the linker cell led collective migration of the male gonad is mechanically very different from the distal tip cell-oriented gonad shaping in hermaphrodites. The linker cell unequivocally moves itself forward, as seen both in mutants with the cells detached and those physically cut off from the rest of the gonad (Chisholm, 1991; Kato et al., 2014; Sternberg and Horvitz, 1988). In both cases, the linker cell continues its normal course of migration. Thus, linker cell migration is a useful genetic model for cell adhesion research, in particular, cell adhesion during migration. We reasoned that linker cell detachment happens in homozygous mutant animals descended from heterozygous mothers that were viable themselves due to 1) The encoded protein was provided maternally earlier in the development, where most other collective migrations occur. 2) The probable intense stress applied to the cells from the tow. This extreme requirement for adhesion might makes this process more susceptible to decreased function of proteins that in other contexts might not display a defect. In this way, the function of broadly utilized proteins might be illuminated in the linker cell context.

### A diverse group of proteins is required to keep the migrating cells together

Mutant animals of three orthologs of four ITFG1 interactors were found to have an apparently identical linker cell detachment phenotype. The phenotype, likely associated with cell adhesion, is also found in the *C. elegans lnkn-1* mutants. Animals carrying mutant alleles of the three orthologs *tat-5*/ATP9A, *ndk-1*/NME1/4, and *apc-2*/ANAPC2, also displayed other phenotypes that we observed in *lnkn-1(lf)* animals, suggesting involvement in similar biological process.

*tat-5*, ortholog of the mammalian ATP9A and ATP9B genes, belong to the eukaryotic P4-type ATPases family of proteins (Lyssenko et al., 2008). These flippase use energy from ATP hydrolysis to drive the active transport of phospholipids between the two leaflets of the membrane (Andersen et al., 2016). The *C. elegans* genome encodes six members of the family, of which *tat-5* is the only one that is essential (Lyssenko et al., 2008). Previous studies suggest that TAT-5 maintains phosphatidylethanolamine (PE) asymmetry in the membrane and suppresses the budding of extracellular vesicles, and loss of *tat-5* results in abnormal embryonic cell shape, which can be argued to result from reduced cell adhesion (Beer et al., 2018; Wehman et al., 2011).

NDK-1 is the sole group I member of the NME proteins in *C. elegans* (Masoudi et al., 2013). Also known as NM23 (non-metastatic 23) / NDPK nucleoside diphosphate kinase, the family is broadly conserved in evolution (Bilitou et al., 2009; Cetkovic et al., 2015). The first member of the family, a group I member, NME1, was the first identified metastasis suppressor gene (Steeg et al., 1988). Two group I NME proteins – NME1 and NME 4 – were enriched in both of the ITFG1-pulldown assays (Table S1-3). Studies of the group in cell lines and model organisms suggest its involvement in developmental processes, including migration and cell proliferation (Matyasi et al., 2020). Notably, NME1 is argued to locally produce GTP for use by GTPases such as dynamin (Farkas et al., 2019). Previous studies of NDK-1 in *C. elegans* suggest its involvement in the Ras/MAPK signaling, apoptosis, and the shaping of the hermaphroditic gonad (Fancsalszky et al., 2014; Farkas et al., 2018; Farkas et al., 2019; Masoudi et al., 2013; Tran et al., 2019).

*apc-2* encodes the worm ortholog of human ANAPC2 (anaphase promoting complex subunit 2). The anaphase promoting complex (APC), also known as cyclosome, is an E3 ubiquitin ligase that is essential for the progression of eukaryotic cell cycles (Peters, 2006). It is thought that the activities of APC can affect gene expression, at least at the chromatin level (Bodrug et al., 2021).

How might these three very different proteins be involved in holding the migrating cells together? Kato et al. (2014) suggest that LNKN-1, with its extracellular domain containing motifs with similarities to that of integrin, might be directly involved in physical adhesion. Based on that model, the intracellular domain of LNKN-1 interacts with RUVB-1, RUVB-2, and alpha-tubulin, connecting the junction to the microtubule cytoskeleton. This hypothesis was supported through protein interactions of their human counterpart in cultured cells, which we again validated in this study, and of colocalization in the worm gonad. Previous studies find the TAT-5 to be associated with cell adhesion (Wehman et al., 2011), presumably depending on its flippase activity that can locally modify membrane composition. It is possible that LNKN-1 and TAT-5 are closely associated at the adhesion junction and reinforce the assembly of a complex promoting strong adhesion. As a nucleoside diphosphate kinase capable of generating GTP from GMP, it is plausible that NDK-1 is recruited by LNKN-1 or its associated protein to provide local activation, perhaps for a yet to be identified G protein that functions locally to promote cell adhesion. Based on the cell detachment phenotype, a few other proteins were also identified, although it is not known whether they are directly involved since no interaction was found between their orthologs and that of ITFG1. Those proteins include EGL-5 (Chisholm, 1991; Schwarz et al., 2012), which, as a homeobox transcription factor, is likely to be indirectly involved. Other proteins are orthologs of each of the SMC (Structural maintenance of chromosomes) proteins 1-4 (Schwarz et al., 2012), which are the core parts of the cohesin and condensin complexes (Hirano, 2006). The addition of a component of the anaphase promoting complex, APC-2, raises the question of why the LINKIN interactome includes significant number of chromosomes associated proteins. A default explanation is that this is due to an indirect effect of cell cycle events on the post-mitotic linker cell. Another hypothesis is that chromosome biology plays a structural role. An insightful review by Bustin and Misteli (2016) argues that chromosomes, as large entities in the cell, have other roles in cell processes unrelated to genome integrity and gene expression. We propose that the linker cell chromosomes help anchor the cytoskeleton and membrane adhesion molecules, making the nucleus a huge anchor that can withstand the force of collective cell migration (Fig. 6).

**Fig. 6.**
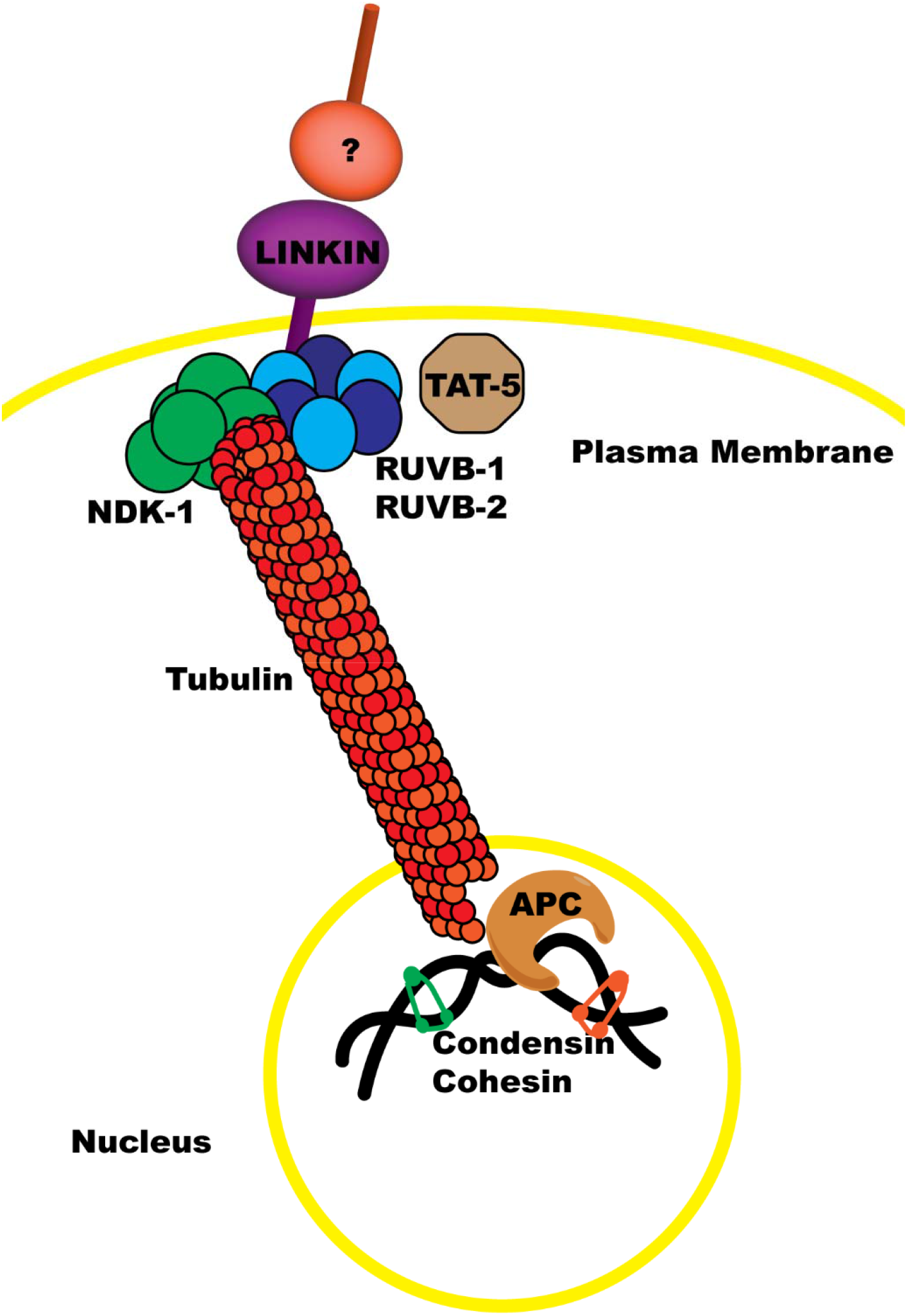
Model for LINKIN-dependent cell adhesion during cell migration. We hypothesize that LNKN-1 promotes cell adhesion through the interactions of its extracellular domain with other cells or extracellular matrix and through its intracellular domain by interacting with the microtubule cytoskeleton, which connects it to the chromosome. This allows the use of the nucleus as a huge anchor that can withstand the force of collective cell migration. RUVB-1/2 interact with LNKN-1 at the plasm membrane, and the microtubule cytoskeleton interacts with the chromosome potentially through the anaphase-promoting complex. NDK-1 and TAT-5 also promote cell adhesion and are likely associated with LNKN-1 at the plasma membrane, while condensin and cohesin likely promote cell adhesion through their interaction with chromosomes.

## Materials and Methods

### Knowledgebases

We used WormBase (Davis et al., 2022), the Alliance of Genome Resources (Alliance of Genome Resources, 2022), STRING (Szklarczyk et al., 2023), and GeneMania (Franz et al., 2018) in the design and interpretation of experiments.

### Plasmid construction and cell lines

The full-length ITFG1 with or without a C-terminal Myc tag (OriGene Technologies, Inc., RC204773) was cloned in the expression vector pcDNA3.3 or pRRL.sin.cPPT.SFFV/IRES-neo.WPRE. The full-length RUVBL1 with N-terminal 3×FLAG tag (pCDNA-3xFLAG-Pontin) was obtained from Addgene (51635). All cell lines were grown in DMEM (Sigma-Aldrich) supplemented with 10% fetal bovine serum (FBS, Atlanta Biologicals) plus 100 U/mL penicillin and 100 µg/mL streptomycin (Lonza) and maintained at 37°C with 5% CO_2._ Transfection was performed using BioT (Bioland Scientific LLC). MDA-MB-231 cells were transiently transfected with pcDNA3.3 expressing ITFG1-Myc or ITFG1. MDA-MB-231 stably expressing ITFG1-Myc were generated by transduction with pseudotyped lentiviral vector produced by transient co-transfection of 293T cells with pRRL.sin.cPPT.SFFV/IRES-neo.WPRE expressing ITFG-Myc plasmid and lentivirus packaging plasmids (pHDM-G, CAG4-RTR2 and CAGGHIVgpco). Stable cell lines were selected in the presence of 5 mg/mL G418 24 h post-lentivirus infection. MDA-MB-231 cells stably expressing ITFG1-Myc were transfected with pCDNA-3xFLAG-Pontin for transient expression of FLAG-RUVBL1.

### Immunofluorescence staining of ITFG1 expression in MDA-MB-231 cells

Cells were seeded in a 96-well plate, which was coated with 20 mg/mL fibronectin (Sigma-Aldrich, F1141). On the next day, cells were fixed in ice-cold 4% paraformaldehyde at room temperature for 5 min. After blocking with 10% FBS plus 0.2% Triton X-100 at room temperature for 1 h, cells were overnight incubated with Rabbit anti-ITFG1 antibody (Invitrogen, PA5-54067) at 4°C, followed by Alexa FluorTm 594 donkey anti-rabbit IgG (H+L) (Invitrogen, A21207) at room temperature for 1.5 h. Cells were counter stained with Hoechst 33342 (Thermo Scientific, 62249) and visualized with an ImageXpress® Confocal HT.ai High-Content Imaging System (Molecular Devices).

### Western blot assay

Cells were pelleted and washed three times by PBS. Fractionation of subcellular proteins were collected using Subcellar Protein Fractionation Kit (Thermo Scientific, 78840). Cell lysates were prepared in lysis buffer (20 mM HEPES pH 7.5, 100 mM NaCl, 0.2% n-dodeyl-D-Maltoside, 1 mM N-Ethylmaleimide, 1% Triton-100, 0.02 mM MG132, and Pierce protease inhibitors (Thermo Scientific, A32965). Protein samples were loaded on 4-20% SDS-PAGE gel (Bio-Rad) and transferred to a nitrocellulose membrane (Bio-Rad). The following primary antibodies were used for protein blotting: mouse anti-c-Myc antibody (Sigma-Aldrich, 05-724MG), mouse anti-Flag antibody (Sigma-Aldrich, F3165), Rabbit anti-ITFG1 antibody (ITFG1_Ab1) (Invitrogen, PA5-54067), mouse anti-ITFG1 antibody (ITFG1_Ab2) (R&D Systems, MAB89001), Rabbit anti-RUVBL1 antibody (Proteintech, 10210-2-AP), Rabbit anti-GAPDH antibody (Cell Signaling Technology, 2118). Signals were developed with HRP-labeled secondary antibodies (Bio-Rad). Blots were developed using Immobilon Western Chemiluminescent HRP Substrate (Millipore) and visualized using ChemiDoc MP Imaging System (Bio-Rad).

### Flow cytometry

Surface expression of ITFG1 on cells was examined by staining the cells with mouse anti-ITFG1 antibody (R&D Systems, MAB89001) at 4°C for 1 hour, followed by Alexa Fluor 488 donkey anti-mouse IgG (H+L) (Invitrogen, A21202). The surface florescence intensity of viable cells was measured by S3e Cell Sorter (Bio-Rad).

### Immunoprecipitation and sample preparation for LC-MS/MS

Each immunoprecipitation was performed in triplicate. Cell lysates were prepared in lysis buffer. Protein concentration was determined using the Bradford protein assay (Bio-Rad). Equivalent amounts of protein (∼2 mg) were used for immunoprecipitation with Pierce anti-c-Myc Agarose (Thermo Scientific, 20169), anti-Flag Affinity Gel (Bimake, B23101), or anti-ITFG1 antibody-conjugated agarose. Two anti-ITFG1 antibodies were covalently conjugated on agarose beads using Pierce Direct IP Kit (Thermo Scientific, 26148), respectively. Immunoprecipitation was performed at 4°C for 2 h in micro-spin column (Thermo Scientific, 89879), and the beads were washed three times with lysis buffer and two times with Mass Spec buffer (0.1 M Tris-HCl in mass grade water, pH 8.5). Proteins were eluted with 10 M urea solution and prepared for LC-MS/MS analysis using EasyPep Mini MS Sample Prep Kit (Thermo Scientific, A40006). The dried peptides were dissolved with 0.1% formic acid.

### LC-MS/MS and protein identification

LC-MS/MS experiments were performed using EASY-nLC1000 (Thermo Fisher Scientific) connected to an Orbitrap Eclipse Tribrid mass spectrometer (Thermo Fisher Scientific). The detailed information for LC/MS processing and data analysis was described previously (Cheng et al., 2021). System control and data collection were performed by Xcalibur v.4.0. Proteomic analysis was performed with the Proteome Discoverer 2.4 (Thermo Scientific) using the SequestHT search algorithm with Percolator validation. The fold changes of each protein expression were compared between ITFG-Myc-expressing cells and non-expressing control cells. We used 200-fold change (log2 FC = 7.64) when the proteins were only detected in ITFG-Myc-expressing cells but none in non-expressing control cells. A set of proteins = upregulated in ITFG-Myc-expressing cells compared to non-expressing control cells in at least two immunoprecipitates were identified (log2 FC > 0.6), in which proteins with higher upregulated (log2 FC > 2) were selected for overlap analysis. Venn plots were generated with FunRich 3.1.4 (Fonseka et al., 2021; Pathan et al., 2015; Pathan et al., 2017). The resulting list of overlapped proteins was subjected to Reactome pathway and Gene Ontology (GO) biological process enrichment analysis using Metascape (Zhou et al., 2019). The RAW data have been deposited to the ProteomeXchange Consortium via the PRIDE partner repository with the dataset identifier PXD044339, user name, and password: GrISpXFX.

### Nematode genetics and general methods

The strains of *C. elegans* were maintained using standard methods similar to what was described by Brenner (1974). Briefly, worms were cultured on Nematode Growth Medium (NGM) dishes seeded with a lawn of *Escherichia coli strain* OP50 at 20°C. All strains were derived from the wild-type reference strain Bristol N2 (Brenner, 1974). Alleles and transgenes used in this study were: LG I: *src-1(cj293)* (Bei et al., 2002), *dpy-5(e61), unc-13(e450)* (Brenner, 1974), *unc-94(ok1210), cdc-25.1(ok1888), ndk-1(ok314)* (Consortium, 2012), *feh-1(gb561)* (Zambrano et al., 2002), *tat-5(tm1741)*, *tmC18[dpy-5(tmIs1200)]*, *tmC18[dpy-5(tmIs1236)], tmC27[unc-75(tmIs1239)]* (Dejima et al., 2018), *let-611(h826)*; LG II: *syIs128* (Kato and Sternberg, 2009); LG III: *apc-2(ok1657)* (Consortium, 2012), *oxTi719*, *oxTi956* (Frokjaer-Jensen et al., 2014), *lnkn-1(sy1596), mvk-1(tm6628), trd-1(tm2764), qC1 [dpy-19(e1259) glp-1(q339)] nIs189, sC1(s2023)[dpy-1(s2170)umnIs21]*; LG IV: *him-8 (Hodgkin et al., 1979), rad-51(ok2218)* (Consortium, 2012), *rod-1(tm6186), tmC5[F36H1.3(tmIs1220)]* (Dejima et al., 2018); LG V: *him-5 (Hodgkin et al., 1979)*, *cdc-25.2(ok597), C10B5.1(ok3270), hsp-90(ok1333), lin-40(gk255), lin-40(ok905), lin-40(ok906)* (Consortium, 2012)*, tmC3[egl-9(tmIs1228)], tmC3[egl-9(tmIs1230)], tmC12[egl-9(tmIs1197)], tmC16[unc-60(tmIs1210)], tmC16[unc-60(tmIs1237)]* (Dejima et al., 2018); LG X: *vps-41(ok3433)* (Consortium, 2012)*, tmC30[ubc-17(tmIs1243)], tmC30[ubc-17(tmIs1247)]* (Dejima et al., 2018).

### Generation of *lnkn-1(sy1596)* loss-of-function allele

The *lnkn-1(sy1596)* knock-out animal was generated by CRISPR/Cas9, using the STOP-IN cassette strategy as described in Wang et al. (2018). The 43bp knock-in cassette (GGGAAGTTTGTCCAGAGCAGAGGTGACTAAGTGATAAgctagc) was inserted near the 5’ end of the coding region between CTTGGAAAAGTATGTGCATTTGGAGATTTCAATGCA and GATCGGAATACTGATATTCTGGTTTTTGCGAATG. The *lnkn-1(sy1593)* (III:-0.01) mutation was maintained *in trans* by two inserted fluorescent marker *oxTi719[eft-3p::tdTomato::H2B]* (III:-0.26) and *oxTi956[eft-3p::GFP::2xNLS::tbb-2]* (III: 0.03) (Frokjaer-Jensen et al., 2014).

### Balancer strain construction

We obtained alleles for candidate genes based on: (1) the existence of an available allele; (2) a judgment based on available information from Worm Base https://wormbase.org (Davis et al., 2022); Caenorhabditis Genetics Center https://cgc.umn.edu; and in NBRP of Japan https://nbrp.jp/en/resource/c-elegans-en, in whether the allele was described as either lethal or sterile; and (3) whether the allele was available without other mutations in *cis*. We then rebalanced many of the obtained alleles with balancers (Dejima et al., 2018) that are intrachromosomal and fluorescently labeled (Table S5).

### Microscopy

Images were acquired with a Zeiss Imager Z2 microscope equipped with an Apotome 2 and Axiocam 506 mono using Zen 2 Blue software. Worms were immobilized with levamisole and mounted on 5% agarose pads on microscope slides for observation.

### Development and fertility assay of hermaphrodite worms

All homozygous hermaphrodites of the lethal or sterile mutations used in this study descended from balanced heterozygotic mothers. homozygous animals were identified through the lack of fluorescently labeled balancers. In Fig. 3C, mutations that prevented worms from adulthood were excluded from further analysis. Phenotype 2 was defined by having F_1_ progenies that hatch. Phenotype 3 was defined by the observation of embryos/ embryo-like objects or oocytes on the NGM plates in which the worm was cultured. Some worms that very rarely (*unc-94*) or never (like *rod-1*) lay anything were observed to possess embryos and oocytes inside the gonad. In the maternal effect experiments (Fig. 3B-C phenotype 4,5), homozygous hermaphrodites were mated with *him-5(e1490)* males carrying fluorescent genetic markers *oxTi719* [*eft-3p::tdTomato::H2B*] and *oxTi956[eft-3p::GFP::2xNLS::tbb-2]*. Due to large differences in brightness, only *oxTi719* was assayed for the experiment. Phenotype 4 was determined by whether the male-derived *oxTi719* marker was observed in F1 embryos/animals, as seen in Fig. 3B-Phenotype 5 was based on whether those F1 (as in phenotype 4) hatch.

### Assessment and measurement of the gonad phenotype in hermaphrodites

The measurement and analysis of the gonad phenotype in hermaphrodites were performed as described in Fig. 4C. The measurement was done using the ImageJ software (NIH) with homozygous hermaphrodites two days past L4. The approximated ventral-proximal gonad arm lengths were measured by drawing a line through the midline of the worm as the distance from the vulval to the ventral-dorsal turn of the U-shaped gonad arm. The approximated dorsal-distal gonad arm lengths were measured similarly, as the distance from the distal end of the gonad to the ventral-dorsal turn. The Centripetal migration (%) was calculated by dividing the value of dorsal-distal gonad arm length by the value of ventral-proximal gonad arm length and multiplying by 100%. Due to the positioning of the worms on the microscope slide, not all worms were measured for both arms of the gonad. Pathfinding or other gonadal development defects were observed in some of the mutants, and animals exhibiting those phenotypes were not included in this examination. All comparison in Fig. 4D was to the value of the wild-type, using two-tailed, unpaired Student’s *t*-test.

### Assessment of male gonad development and linker cell migration analysis

For Fig. 5, all homozygous males descended from balanced heterozygotic mothers, and all carry the transgene *syIs128 [lag-2p::YFP]* (Kato and Sternberg, 2009), which labels the cytoplasm of the linker cell. In addition, all strains carry either *him-5(e1490)* or *him-8(e1489)* (Hodgkin et al., 1979) to increase male occurrence. The male *apc-2(ok1657)* homozygous males have other clear tail morphological defects post-L4, but the linker cell detachment precedes that development. In Fig. 6B-E, the linker cell was identified through both the cell morphology under DIC and the transgene *syIs128*.

### RNA extraction and qPCR analysis

Total cell RNA was extracted from cells using the MagMAX *mir*Vana Total RNA Isolation Kit (Thermo Scientific, A27828) with KingFisher Duo Prime Purification System (Thermo Scientific). Total RNA was converted to cDNA using High-Capacity cDNA Reverse Transcription Kit (Thermo Scientific, 4368813). Quantitative PCR (qPCR) was performed on the QuantStudio™ 5 Real-Time PCR System (Thermo Scientific) using PowerUp SYBR Green Master Mix (Applied Biosystems, A25741). Primer sequences of ITFG1 used in qPCR: TGGGAGCTGACAGACCTAAA and GCAGTAAGCAGAACAATATTACTTGG. ITFG1 RNA level was normalized to GAPDH as a reference gene.

### In-gel digestion

Cell lysates from ITFG1-Myc-expressing MDA-MB-231 and non-expressing control MDA-MB-231 cells were separated in a 4-20% SDS-PAGE gel (Bio-Rad). The gels were stained with Imperial Protein Stain (Thermo Scientific, 24615). Gel lanes in the molecular weight range between 150 and 250 kDa, 75 and 150 kDa, and 37 and 75 kDa were removed and further excised into small pieces 1.5 ml tube followed by in-gel digested using Tryptic Digestion Kit (Thermo Scientific, 89871) according to the manufacturer’s instructions. After digestion, digested mixtures were dried using vacuum centrifugation and dissolved with 0.1% formic acid for LC-MS/MS analysis.

## Acknowledgements

We thank members of our laboratory especially Hillel Schwartz and Nick Markarian for discussions. Mutant alleles – *mvk-1(tm6628)*, *trd-1(tm2764), vha-19(tm2225), prx-3(tm6469),* and *rod-1(tm6186)* – were kindly provided by the MITANI Lab through the National Bio-Resource Project of the MEXT, Japan. Some strains were obtained from the Caenorhabditis Genetics Center (CGC), which is funded by the NIH Office of Research Infrastructure Programs (P40 OD010440). We thank the Chan laboratory for use of their flow cytometer. This work was also facilitated by WormBase, a knowledgebase for nematode research; and by the Alliance of Genome Resources, a research platform that facilitates cross species research. The Proteome Exploration Laboratory (PEL) is supported by the Beckman Institute at Caltech. This research was supported by NIH R01HD086596 (PWS and TFC), R01HD091327 (PWS), and R24 0D023041 (PWS).

---

**Fig. S1.**
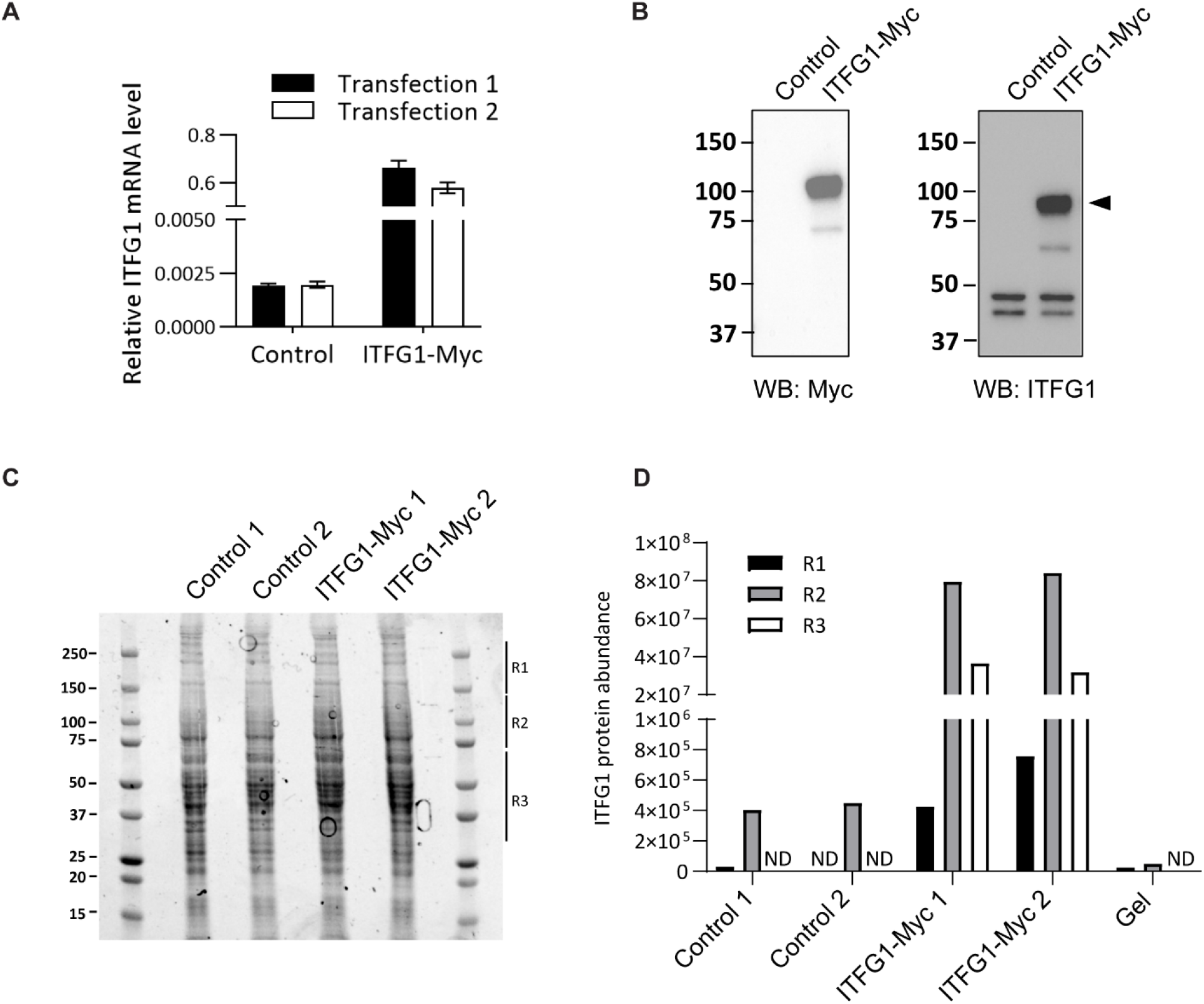
ITFG1 identification in ITFG1-Myc-expressing MDA-MB-231 cells. ITFG1-Myc-expressing MDA-MB-231 cells and non-expressing control cells were harvested at 48 h post transient transfection for determining ITFG1 expression. (A) ITFG1 RNA in cell lysates was quantified by real-time PCR. Data show ITFG1 RNA levels relative to GAPDH. Two individual transfections were analyzed. Error bars represent SD (n = 3). (B) Representative western blot images showing the expression of ITFG1 and GAPDH (loading control) in cell lysates. ITFG1 expression was probed with anti-Myc (left) or anti-ITFG1 antibodies (right). (C) Coomassie blue staining of SDS-PAGE for Myc-immunoprecipitates from ITFG1-Myc expressing and non-expressing control MDA-MB-231 cells. Gel lanes in the molecular weight range between 150 and 250 kDa (R1), 75 and 150 kDa (R2), and 37 and 75 kDa (R3) were removed followed by in-gel digested for LC/MS-MS analysis. (D) ITFG1 protein abundance in each region identified by LC/MS-MS.

**Fig. S2.**
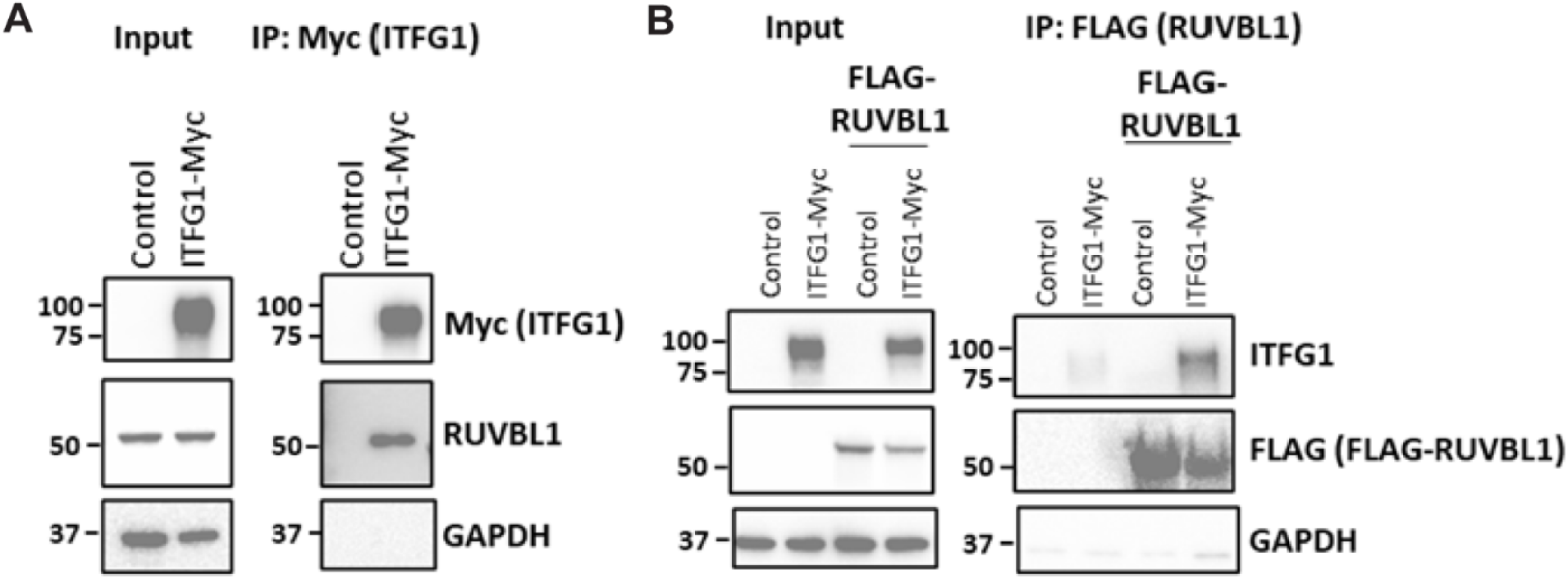
ITFG1-RUVBL1 interaction in ITFG1-expressing MDA-MB-231 cells. (A) Western blot assay identified ITFG1 and RUVBL1 in Myc-immunoprecipitants in ITFG1-Myc transient expressing cells using anti-Myc and anti-RUVBL1 antibodies, respectively. (B) Western blot assay identified ITFG1 and RUVBL1 in FLAG-immunoprecipitants in FLAG-RUVBL1 transient expressing cells using anti-ITFG1 and anti-FLAG antibodies, respectively. Input from cells was used as the positive control. GAPDH was used as the loading control for input and negative control for immunoprecipitation. IP, immunoprecipitation.

