## Supplementary material for "LINKIN-associated proteins necessary for tissue integrity during collective cell migration": Table S4

| **Table S4 Worm orthologs of selected ITFG1 interactors** | | | | |
| --- | --- | --- | --- | --- |
| **Accession** | **Gene Symbol** |  | **Re-Balanced** | **Analyzed** |
| P54652 | HSPA2 | *hsp-1* | Y | Y |
| G5E9A6 | USP11 | H34C03.2 |  |  |
| Q8IWQ3 | BRSK2 | *sad-1* |  |  |
| Q96EB6 | SIRT1 | *sir-2.1* |  |  |
| F8W8A6 |  | *hach-1* |  |  |
| P36896 | ACVR1B | *sma-6* |  |  |
| P07948 | LYN | *src-1* | Y |  |
| Q96DA2 | RAB39B | *rab-39* |  |  |
| Q9BVT8 | TMUB1 | B0303.4 |  |  |
| Q969Y2 | GTPBP3 | *mtcu-1* |  |  |
| P42574 | CASP3 | *ced-3* |  |  |
| P42574 | CASP3 | *csp-1* |  |  |
| Q6JQN1 | ACAD10 | *acds-10* |  |  |
| P98172 | EFNB1 | *efn-2* |  |  |
| P98172 | EFNB1 | *efn-3* |  |  |
| P98172 | EFNB1 | *vab-2* |  |  |
| A0A0A0MTL5 |  | *skpt-1* |  |  |
| Q9Y535 | POLR3H | *rpc-25* |  |  |
| P01024 | C3 | *tep-1* |  |  |
| Q9Y2Y1 | POLR3K | *rpc-11* |  |  |
| F5H8H2 | MVK | *mvk-1* | Y | Y |
| Q9H871 | RMND5A | *gid-2* |  |  |
| P30307 | CDC25C | *cdc-25.1* | Y | Y |
| P30307 | CDC25C | *cdc-25.2* | Y | Y |
| P0CG13 | CHTF8 | *ctf-8* |  |  |
| O94776 | MTA2 | *lin-40* | Y | Y |
| Q9BVN2 | RUSC1 | *unc-14* |  |  |
| P49754 | VPS41 | *vps-41* | Y | Y |
| B5MCD7 | SYNGR1 | *sng-1* |  |  |
| P50748 | KNTC1 | *rod-1* | Y | Y |
| Q147X3 | NAA30 | *natc-2* |  |  |
| Q9UJX6 | ANAPC2 | *apc-2* | Y | Y |
| O60783 | MRPS14 | *mrps-14* |  |  |
| Q8WWC4 | C2orf47;  MAIP1 | Y62E10A.20 |  |  |
| P52824 | DGKQ | *dgk-1* |  |  |
| Q4TT34 | NME4 | *ndk-1* | Y | Y |
| O95905 | ECD | F19C6.2 | Y | Y |
| Q58FF6 | HSP90AB4P | *hsp-90* | Y | Y |
| P56589 | PEX3 | *prx-3* | Y | Y |
| A0A075B7G8 |  | *feh-1* | Y | Y |
| Q06609 | RAD51 | *rad-51* | Y | Y |
| Q9Y4U1 | MMACHC | *cblc-1* |  |  |
| P56282 | POLE2 | *pole-2* |  |  |
| Q9H4L7 | SMARCAD1 | M03C11.8 |  |  |
| Q9UJX4 | ANAPC5 | *gfi-3* |  |  |
| Q9UJX4 | ANAPC5 | *such-1* |  |  |
| Q5R372 | RABGAP1L | *tbc-11* |  |  |
| Q96SZ6 | CDK5RAP1 | F25B5.5 |  |  |
| O75419 | CDC45 | *evl-18* |  |  |
| Q8TB96 | ITFG1 | *lnkn-1* |  |  |
| Q9Y4R8 | TELO2 | *clk-2* |  |  |
| Q15904 | ATP6AP1 | *vha-19* | Y | Y |
| Q9Y287 | ITM2B | C25F6.7 |  |  |
| Q99956 | DUSP9 | *lip-1* |  |  |
| Q96L35 | EPHB4 | *vab-1* |  |  |
| Q9NQE9 | HINT3 | *hint-3* |  |  |
| Q9Y230 | RUVBL2 | *ruvb-2* |  |  |
| Q9NVI1 | FANCI | *fnci-1* |  |  |
| Q9BYB4 | GNB1L | C10B5.1 | Y | Y |
| Q9NVH6 | TMLHE | *gbh-2* |  |  |
| Q9BZX2 | UCK2 | B0001.4 |  |  |
| Q9P2R3 | ANKFY1 | F22G12.4 |  |  |
| C9JJ19 | MRPS34 | mrps-34 |  |  |
| O75110 | ATP9A | *tat-5* | Y | Y |
| Q9NX20 | MRPL16 | *mrpl-16* |  |  |
| P52564 | MAP2K6 | *sek-1* |  |  |
| P0DN79 | CBS; LOC102724560; CBSL | *cbs-2* |  |  |
| Q9BYD2 | MRPL9 | *mrpl-9* |  |  |
| P10586 | PTPRF | *ptp-3* |  |  |
| Q9BUI4 | POLR3C;  LOC101060460 | *let-611* | Y |  |
| Q6NZY4 | ZCCHC8 | ZK632.11 |  |  |
| Q6NZY4 | ZCCHC8 | Y34D9A.7 |  |  |
| Q9Y673 | ALG5 | *algn-5* |  |  |
| Q6P996 | PDXDC1;  LOC102724985 | C14H10.3 |  |  |
| P15531 | NME1 | *ndk-1** | Y | Y |
| O75600 | GCAT | T25B9.1 |  |  |
| P08243 | ASNS | *asns-1* |  |  |
| P08243 | ASNS | *asns-2* |  |  |
| Q0VAK6 | LMOD3 | *tmd-2* |  |  |
| Q0VAK6 | LMOD3 | *unc-94* | Y | Y |
| Q6P3X3 | TTC27 | *trd-1* | Y | Y |
| Q7Z7H8 | MRPL10 | *mrpl-10* |  |  |
| Q9Y265 | RUVBL1 | *ruvb-1* |  |  |
| O43175 | PHGDH | C31C9.2 |  |  |
| Q13405 | MRPL49 | *mrpl-49* |  |  |
