## Supplementary material for "LINKIN-associated proteins necessary for tissue integrity during collective cell migration": Table S5

| **Table S5 Balanced strain constructed for this study** | | | |
| --- | --- | --- | --- |
| **Genes** | **Allele source** | **Re-balanced Strain**  **Genotype** | **LC labeled Strain**  **Genotype** |
| *lnkn-1* | PS8994 | **PS9073***  *lnkn-1(sy1596)/oxTi719[eft-3p::tdTomato::H2B] oxTi956[eft-3p::GFP::2xNLS::tbb-2] III* | **PS9824**  *syIs128 II; lnkn-1(sy1596)/ oxTi719[eft-3p::tdTomato::H2B] oxTi956[eft-3p::GFP::2xNLS::tbb-2] III; him-5 (e1490) V* |
| *src-1* | HR1275 | **PS9411**  *src-1(cj293) dpy-5(e61)/ tmC18[dpy-5(tmIs1200)]* |  |
| *unc-94* | VC828 | **PS9732**  *unc-94(ok1210)/tmC18[dpy-5(tmIs1236)] I* | **PS9825**  *unc-94(ok1210)/tmC18[dpy-5(tmIs1200)] I; syIs128 II; him-5(e1490) V* |
| *let-611* | KR1482 | **PS9731**  *let-611(h826) dpy-5(e61) unc-13(e450)/tmC27[unc-75(tmIs1239)] I* |  |
| *cdc-25.1* | VC1391 | **PS9394**  *cdc-25.1(ok1888)/tmC18[dpy-5(tmIs1236)] I* | **PS9826**  *cdc-25.1(ok1888)/tmC18[dpy-5(tmIs1200)] I; syIs128 II; him-5(e1490) V* |
| *tat-5* | FT207 | **PS9733**  *tat-5(tm1741)/tmC18[dpy-5(tmIs1236)] I* | **PS9804**  *tat-5(tm1741)/tmC18[dpy-5(tmIs1200)] I; syIs128 II; him-5(e1490) V* |
| *ndk-1* | EJ810 | **PS9406#**  *ndk-1(ok314)/tmC18[dpy-5(tmIs1200)] I; him-8(e1489)/+ IV* | **PS9820**  *ndk-1(ok314)/tmC18[dpy-5(tmIs1236)] I; syIs128 II; him-5(e1490) V* |
| *feh-1* | NA653 | **PS9408**  *feh-1(gb561)/ sC1(s2023)[dpy-1(s2170)umnIs21] III* |  |
| *mvk-1* | FX17538 | **PS9738**  *mvk-1(tm6628)/ qC1 [dpy-19(e1259) glp-1(q339)] nIs189 III* |  |
| *trd-1* | FX18585 | **PS9739**  *trd-1(tm2764)/ qC1 [dpy-19(e1259) glp-1(q339)] nIs189 III* |  |
| *apc-2* | VC1203 | **PS9735**  *apc-2(ok1657)/ qC1 [dpy-19(e1259) glp-1(q339)] nIs189 III* | **PS9803**  *syIs128 II; apc-2(ok1657)/ qC1 [dpy-19(e1259) glp-1(q339)] nIs189 III; him-5(e1490) V* |
| *rod-1* | FX17541 | **PS9740**  *rod-1(tm6186)/ tmC5[F36H1.3(tmIs1220)] IV* | **PS9822**  *syIs128 II; rod-1(tm6186)/tmC5[F36H1.3(tmIs1220)] IV; him-5(e1490) V* |
| *rad-51* | VC1873 | **PS9412**  *rad-51(ok2218)/ tmC5[F36H1.3(tmIs1220)] IV* | **PS9823**  *syIs128 II; rad-51(ok2218)/tmC5[F36H1.3(tmIs1220)] IV; him-5(e1490) V* |
| *cdc-25.2* | YHS25 | **PS9407**  *cdc-25.2(ok597)/tmC16[unc-60(tmIs1237)] V* | **PS9827**  *syIs128 II; cdc-25.2(ok597)/him-8(e1489) IV; tmC16[unc-60(tmIs1210)] V* |
| *lin-40* | VC490 | **PS9410**  *lin-40(gk255)/tmC16[unc-60(tmIs1237)] V* | **PS9828**  *syIs128 II; him-8(e1489) IV; lin-40(gk255)/tmC16[unc-60(tmIs1210)] V* |
|  | VC646 | **PS9736**  *lin-40(ok906)/tmC16[unc-60(tmIs1237)] V* |  |
|  | VC660 | **PS9737**  *lin-40(ok905)/tmC16[unc-60(tmIs1237)] V* |  |
| *C10B5.1* | VC2579 | **PS9734**  *C10B5.1(ok3270)/tmC3[egl-9(tmIs1230)] V* | **PS9821**  *syIs128 II; him-8(e1489) IV; C10B5.1(ok3270)/tmC3[egl-9(tmIs1228)] V* |
| *hsp-90* | VC914 | **PS9405**  *hsp-90(ok1333)/ tmC12[egl-9(tmIs1197)] V* |  |
| *vps-41* | VC2784 | **PS9409**  *vps-41(ok3433)/tmC30[ubc-17(tmIs1247)] X* | **PS9805**  *syIs128 II; him-5(e1490) V; vps-41(ok3433)/tmC30[ubc-17(tmIs1243)] X* |
| Note: The *syIs128* insertion carries the *lag-2p::YFP* marker that labels the cytoplasm of the linker cell. See the Materials and Methods section for more detail.  *: The *lnkn-1(sy1593)*(III:-0.01) KO mutation was maintained by using the two inserted fluorescent marker *oxTi719[eft-3p::tdTomato::H2B]* (III:-0.26) and *oxTi956[eft-3p::GFP::2xNLS::tbb-2]* (III: 0.03). See the Materials and Methods section for more detail.  #: *him-8(e1489)* had not been outcrossed from the strain. | | | |
